# Threatened bat species reduce their activity in presence of traffic noise playback, and may shout louder

**DOI:** 10.1101/2025.11.21.689748

**Authors:** Alisha Hart, Patrick Monari, Kerry Borkin, Kristal E. Cain, Tim Prebble, Ellen Cieraad, David E. Pattemore

## Abstract

Anthropogenic sound is a prevalent pollutant with broad effects on wildlife. As human populations grow, exposure of natural environments to anthropogenic noise will increase. Automobile traffic is a notably pervasive source of noise pollution; transportation networks often cross otherwise relatively undisturbed habitats and the sound they produce is typically chronic and of high intensity. How this noise interferes with the production and reception of animal acoustic signals for communication and navigation remains largely unknown. In particular, bats (order Chiroptera) have received limited attention in anthropogenic noise studies, despite being the second most diverse mammalian group and relying strongly on acoustic signals. We used a ‘phantom road’ paradigm to experimentally isolate and assess the effect of traffic sound playback on the behaviour of the two extant bat species in Aotearoa New Zealand. To begin to understand how bats might behaviourally respond to a noisy environment, we also investigated whether bats’ call sequence and structure differed with and without sound playback. We found an approximate 50% reduction in the mean number of sequences per night of both bat species during sound playback, suggesting bats avoid noisy areas. We also found that calls from the bats that were present were emitted at higher frequencies and increased intensities (loudness) during playback. As avoidance of areas reduces the amount of quality habitat accessible and call modulation can impose metabolic costs, chronic noise may impose fitness costs on declining populations of native bats. With increasing areas of natural habitat exposed to anthropogenic noise, untangling the effects of high sound levels is crucial for conservation of bats and the ecosystem services they provide.

## INTRODUCTION

Anthropogenically generated sound is increasingly recognised as a prevalent pollutant with broad and potentially severe impacts on wildlife (Jerem & Mathews, 2021; Kight & Swaddle, 2011; Shannon et al., 2016). As human populations grow globally, requirements for housing, transportation, and resources increase – driving progressive exposure of natural environments to anthropogenic sound (Jerem & Mathews, 2021; Slabbekoorn & Ripmeester, 2008). Sound from land, air, and water traffic is notably pervasive as transportation networks often cross habitats that may otherwise be relatively undisturbed, and sound from these sources is typically chronic and high-intensity (Potvin, 2017). How this sound interferes with how animals relay and receive acoustic signals that drive their behaviour remains largely unknown.

Many taxa rely on acoustic signals and cues – from both abiotic and biotic sources – to navigate their environment or detect prey, predators, and social partners, and for inter and intra-specific communication (Potvin, 2017). Anthropogenic sound can interfere with the reception and interpretation of acoustic signals in one of three recognised ways (Dominoni et al., 2020; Luo, Siemers, et al., 2015):

1. Masking: when irrelevant noise obscures informative acoustic signals and cues. ‘Masking’ has been used with varying definitions in literature, and is used here where there is spectral and temporal overlap between the irrelevant noise and the informative signal/ cue (e.g., Erbe & Farmer, 2000; Gomes et al., 2016).
2. Distraction/reduced attention: when noise divides the attention of the receiver and decreases their efficiency in singling out important sounds (Allen et al., 2021; Chan et al., 2010).
3. Aversion/misleading: this occurs when the noise is interpreted as a threat, stressor, or annoyance (Luo, Siemers, et al., 2015). This may involve the noise being mistaken for a relevant cue/ signal which is aversive (e.g., a predator, heavy rain) (Geipel et al., 2019).

Animals have two key ways of coping with anthropogenic noise: avoidance and mitigation (Figure 1). The first strategy simply avoids scenarios where noise levels lead to compromised signal or cue detection. Avoidance may be spatial, such as the movement of cod (*Gadus marhua*) away from boat engine noise (Mickle & Higgs, 2018), or temporal, like the shifting of the dawn chorus earlier by some bird species to avoid noisy rush hour traffic (Arroyo-Solís et al., 2013).

**Figure 1.**
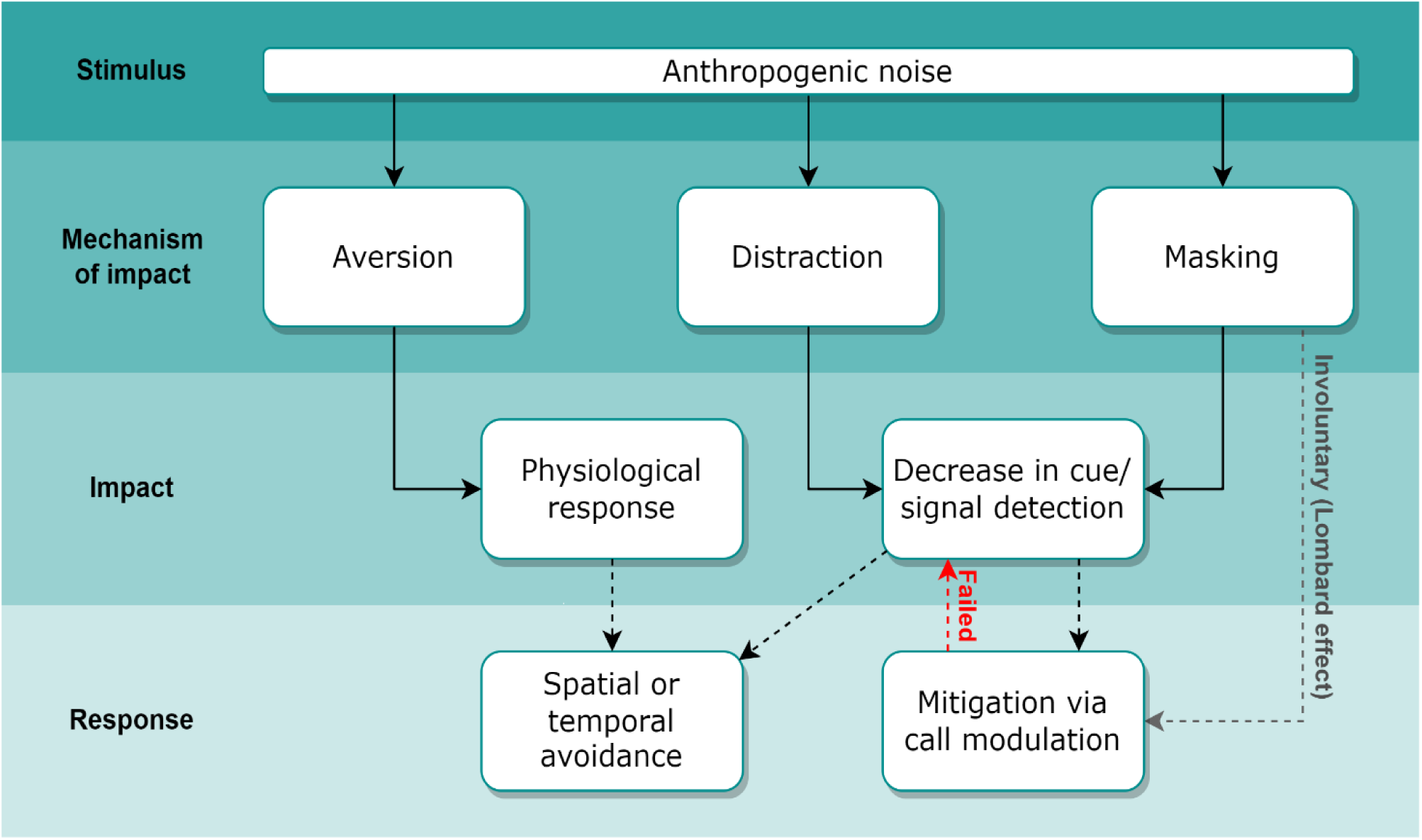
Schematic showing the chain of mechanisms, negative impacts, and immediate behavioural responses currently established for bats exposed to anthropogenic noise. Aversion covers theories such as annoyance or ‘misleading’ where the sound is perceived as a threat. Dotted lines indicate potential responses to impacts, which may not occur.

The second strategy mitigates the masking effect by altering call properties. This may be an immediate, temporary alteration to signals, or long-term adaptation arising from selection of acoustic signalling traits most effective in areas subject to chronic anthropogenic noise (Potvin, 2017). Acoustic signals can be modified in several ways to reduce the effect of masking. These are broadly known as the ‘Lombard effect’, an involuntary increase in vocal effort in response to masking noise, which can include many call characteristic changes including amplitude, frequency, call rate, bandwidth, and duration (Brumm & Zollinger, 2011; Hotchkin & Parks, 2013). Commonly reported ways to avoid overlap with background noise are the increase in signal amplitude (i.e. calling more loudly to improve the signal-to-noise ratio) or a shift in acoustic frequency (i.e. raised pitch to reduce overlap with masking noise) (Brumm & Slabbekoorn, 2005). However, other changes such as increasing call duration or increasing call rate/ decreasing intervals between calls may also make them more detectable, when ‘competing with’ fluctuating or intermittent background noise (Francis, 2015).

While there has been considerable recent research into the effects of anthropogenic sound on wildlife over the last decade, it has been heavily biased towards diurnal avifauna and marine mammals (Jerem & Mathews, 2021). There is a marked lack of research on nocturnal bats (order Chiroptera), which comprise approximately 20% of all extant mammal species (Dutheil et al., 2021), a large portion of which echolocate. Echolocating bats largely vocalise in ultrasonic frequencies (>20 kHz) and rely on acoustic cues and signals for social communication, navigation, and foraging. Anthropogenic sound could interfere with their ability to successfully hear and interpret their own vocalisations and echoes, the calls of conspecifics, activity of prey and predators, and other environmental cues such as running water. With increasing urbanisation and ongoing human population growth (Kundu & Pandey, 2020), there is mounting pressure to understand whether anthropogenic sound – particularly from pervasive sources such as transportation networks – negatively impacts these acoustic-reliant species, so that this impact can be avoided or mitigated if necessary.

Aotearoa New Zealand (henceforth Aotearoa) has two known extant endemic and threatened bats (pekapeka in te reo Māori, the indigenous language; O’Donnell et al., 2023). Habitat loss and disturbance, predation by introduced mammals, and reduced availability of/ competition for roost sites are thought to be the primary factors in observed population and distribution declines (O’Donnell et al., 2018; O’Donnell, 2000). Habitat loss or declines in habitat suitability could occur if bats avoid anthropogenic sound, with potential consequences for populations.

Here we use a ‘phantom road’ paradigm to experimentally assess the effect of traffic sound playback on call sequence detection rates of *Chalinolobus tuberculatus* (long-tailed bat) and *Mystacina tuberculata rhyacobia* (lesser short-tailed bat, hereafter short-tailed bat). The two species occupy different niches. Long-tailed bats are considered edge-adapted, with activity predominantly recorded along forest margins or other edges, where they forage aerially for insect prey. In contrast, short-tailed bats (*Mystacina tuberculata*) are clutter-adapted, most often occupying forest interiors, eating insects and plants both when flying and hunting on the ground. We also investigate whether bat call sequence detection rates and call structure differ with or without traffic sound playback, to begin to understand which mitigation strategies, if any, Aotearoa bat species use to cope with a noisy environment.

If anthropogenic noise from traffic and roads negatively affects a bat’s ability to hear important signals or acts as an aversive stimulus, then we predicted we may see bats avoiding affected areas – both spatially and temporally. If so, then we should see a reduction in bat call sequence detections during playback relative to other times and places without playback, as well as greater reductions closer to the playback source. Further, we predicted that if bats do not avoid the area, they will adjust their vocalisations during playback to improve signal detection, resulting in altered call characteristics such as increased call intensity (loudness) and frequency (pitch).

## MATERIALS AND METHODOLOGY

### Study site

Our experiment took place in Waipapa Ecological Area, part of Pureora Forest Park (latitude −38.47, longitude 175.60); an area of protected land covering 78,000 ha of the central North Island of Aotearoa, and one of the few known locations where both species still coexist.

### Phantom road setup

To isolate and assess the effect of traffic sound playback on bat activity and behaviour, without confounding factors such as light or wildlife-vehicle collisions, we used a ‘phantom road’: a line of speakers playing sound recorded from a highway.

The phantom road consisted of three sets of speakers spaced along a deer fence at the forest edge (Figure 2). The speakers were set on 1 m tall tripods 20 m apart, facing the forest; the outer two speaker sets included subwoofers (as lower frequencies attenuate less with distance). The traffic sound file used in playback was recorded 20 m from the edge of the Petone Motorway (State Highway 2, Wellington, New Zealand), a busy dual carriageway with a 100 km/hr speed limit and an annual average daily traffic of approximately 34 000 vehicles, with 4% heavy vehicles (Waka Kotahi, 2021). Sound levels measured 2 m from speakers during playback were highly variable owing to the nature of traffic noise but ranged between 65-80 dB, compared with average nighttime background noise levels without playback of 36.08 ± 3.74 dB. We provide more details in the Supplementary Information.

**Figure 2.**
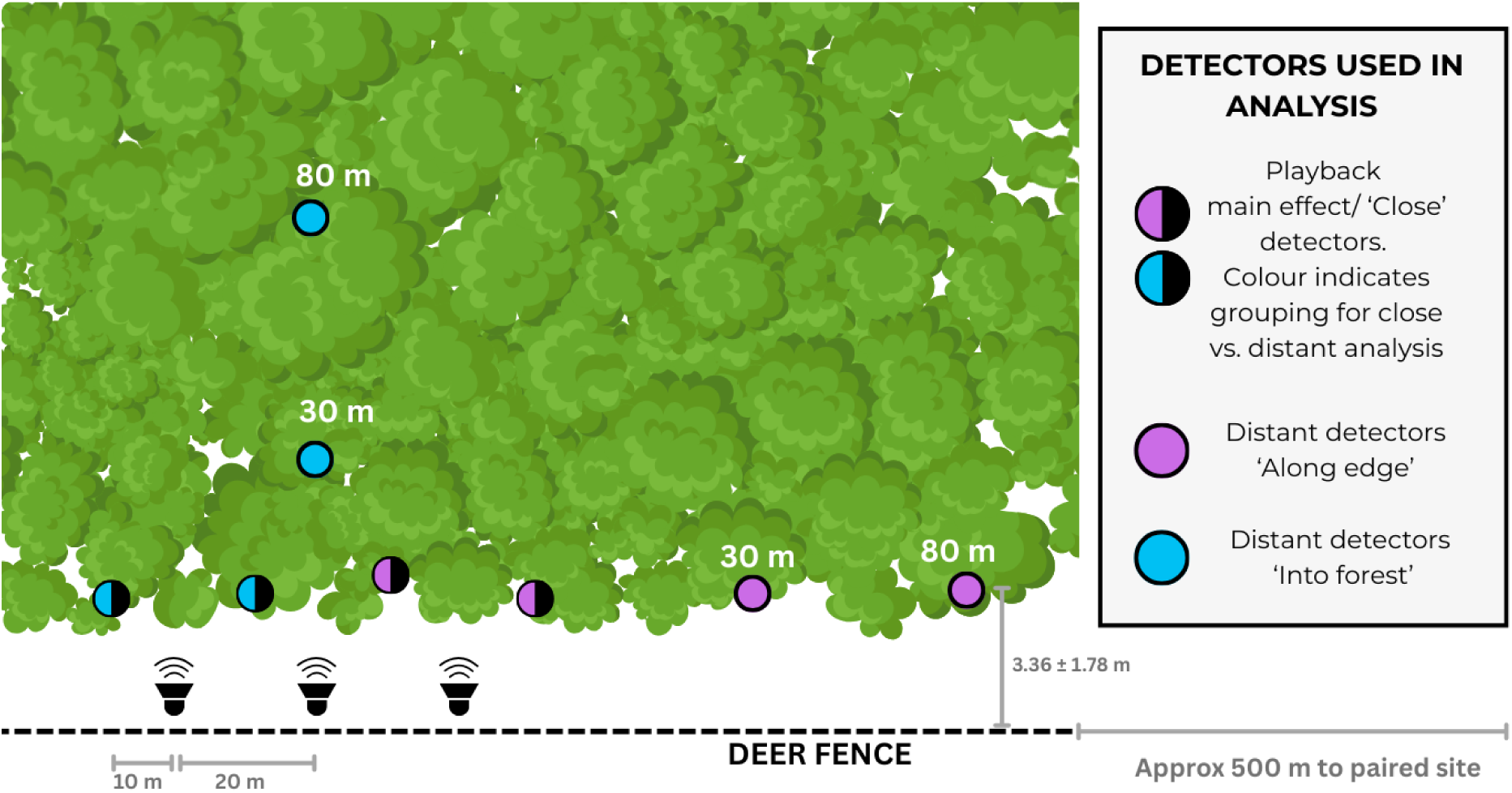
Diagram of speaker and main detector arrangement at each of the four study sites A-D. The three speakers sets are spaced 20 m apart along a deer fence, facing the forest. The outer two speaker sets included the subwoofers. Acoustic bat detectors (circles) were spaced 20 m apart from each other, parallel to the speaker transect, as close as possible to the forest edge, and an average of 3.36 m from the deer fence. Additional detectors were placed at 30 and 80 m into the forest, and along the forest edge towards the paired site. Key shows how distant detectors were grouped for analysis.

### Bat echolocation detection

To record echolocation call sequences of both bat species (henceforth sequences), four acoustic detectors (an omnidirectional frequency compression ultrasound recorder, AR4, Department of Conservation, Wellington, New Zealand) were positioned along the phantom road halfway between each set of speakers and 10 m past either end of each speaker set (Figure 2). Additional detectors were positioned at 30 m and 80 m from the playback speakers, both into the forest and along the forest edge towards the paired site, to assess the effects of traffic playback on sequence detections as a function of distance.

The detectors allowed unobtrusive monitoring by detecting and recording sequences within a detection range of up to 30-50 m (S. Cockburn as cited in D. H. Smith et al., 2020), with each sequence of ultrasonic pulses captured and saved as a spectrogram for later review. Detectors were only 20 m apart within sites, so a degree of overlap in detected bats was expected. Each night, recorders were set to monitor for at least 12.5 hours, starting at least 30 minutes before dusk until at least 30 minutes after sunrise, ensuring sequences would be captured with a buffer either side of dusk and dawn, in case of bats leaving roosts unusually early or returning late in response to the treatment.

To monitor changes in individual call sequences and their characteristics, we used full-spectrum bat detectors (Batlogger A+, Elekon, Luzern, Switzerland) during a subset of nights. This detector type records sounds at their original frequencies and retains a high level of detail of entire calls (Bat Conservation Trust, 2025). At Site A, one detector was placed at Forest-pasture edge adjacent to playback, and one 200 m from the edge in the adjacent open pasture; with one additional detector placed in the adjacent forest interior 50 m from Site B. Detectors were set to record what it considered bat calls from 1 h before sunset to one hour after sunrise each night. Due to limited access to these specialised acoustic recorders, data were captured during 6 nights at the A and B sites, with two nights of playback at site A. The first night of playback coincided with the subset of the full-spectrum bat detector setup, so individual call and call sequence characteristics were only available for one part night and one full night of recording during playback.

### Experimental design

To assess the effect of traffic sound playback on bat call sequence detection rates, we used a multiple before–after control–impact paired series (M-BACIPS) design (Stewart-Oaten et al., 1986), consisting of two paired sites — each containing a control and an impact site — sampled simultaneously on numerous occasions before, during, and after the playback. This allowed us to account for temporal variation (for example caused by weather and seasonal changes). To account for spatial variation, we added a cross-over component, swapping the control and impact sites within each pair.

We selected four sites (A-D) along the park boundary at forest edges where long-tailed and short-tailed bats were known to be present, and that met the following criteria: located along the deer fence that separates the park’s forest edge from the adjacent farmland, with roughly level terrain, at least 250 m and 500 m from the closest public unsealed and sealed road, respectively, and >500 m away from other sites. The four sites were divided into two pairs: pairing A and B on the southern forest edge, and C and D on the eastern edge. All sites had an identical array of detectors (see below).

Following a standard M-BACIPS design, one site in each pair would act as a treatment location (traffic recording playback, see below), while the other served as a control (no playback). One treatment cycle lasted six days, consisting of alternating nights of playback and no-playback until the impact site experienced three playback nights. After one treatment cycle, the control and impact locations were swapped, so that each site was exposed to playback over the study period. The detectors and playback array were then moved to the second site pair, and the experiment was repeated (see Supplementary Materials).

Playback of the looped 53-minute traffic recording was automated, commencing approximately 1 hour before sunset, with a gradual increase in volume over the first 10 minutes to avoid unnecessary startling of nearby wildlife. To avoid potential disturbance of dawn chorus activity of local avifauna, playback stopped approximately 1 hour before dawn. The traffic recording was also played during treatment days from approximately 10:00 to 16:00 to investigate diurnal fauna responses (not detailed in this paper).

Data were collected in Austral summer/autumn, between 4 February and 10 March, 2022. Data collection was scheduled every night unless average wind speeds were expected to exceed 5 m/s, rainfall over 5 mm was predicted, or if the temperature at sunset was below 10°C, as these conditions result in lower bat detection rates (Borkin et al., 2023). The 26 nights of sampling were conducted as planned, with the exception of technical issues resulting in the sound playback stopping early on the first night of playback (Figure S1). This first night of playback was excluded from analyses, as were data from detector 3 at Site C due to a suspected faulty microphone (which caused high interference on the spectrograms and minimal bat detection compared to the other three detectors at Site C).

### Bat call sequence processing

Echolocation spectrograms were processed using AviaNZ software (Avianz.net, Wellington, New Zealand; Marsland et al., 2019) and the ‘NZ Bats_NP’ recogniser, before being manually reviewed by one individual (A. Hart) using the software BatSearch 3.11 (Department of Conservation, Wellington, New Zealand).

Each spectrogram containing a series (sequence) of one or more ultrasonic bat calls (vocalisations) was counted as a single ‘sequence’ for each identified species, or as a sequence for each if vocalisations from both species were present (as in Schamhart et al., 2024). When irregular and/or frequency-shifted pulses in a single spectrogram indicated more than one individual of the same species was vocalising, this was counted as two sequences when there were at least three pulses from each bat (note, a maximum of two sequences were counted per spectrogram, even if three or more bats may have been vocalising). A ‘foraging sequence’ was noted when a spectrogram included a terminal phase or feeding buzz i.e., a rapid call rate over a short period, for example, immediately before bats capturing their target prey; or approach phase, when bats pursue their target and increase pulse repetition rate (Griffin et al., 1960). Ambiguous spectrograms were checked by a second expert (K. Borkin) to establish a consensus, erring on the side of caution, i.e. only clear foraging sequences were noted as such. After removals, the dataset from recorders at the speaker array included approximately 28,500 bat passes, comprised of 95% long-tailed bat passes and 5% short-tailed bat passes.

When investigating the potential effects of playback on call characteristics, we defined a ‘call sequence’ as a spectrogram containing one or more bat calls. One expert manually classified call sequences by species in BatExplorer software (Elekon, Luzern, Switzerland). For call characteristics analyses, we used calls made by single bats which were classified as ‘search phase’ calls - when the pulse repetition of bat calls is relatively low (see supplementary information for more details). Only 67 retained sequences were from lesser short-tailed bats, with 20 of those at the playback site. We do not report on these sparse data here. Of long-tailed bat sequences, we report on results from call characteristics at site A where playback occurred, resulting in 1.5 nights with playback (due to playback cutting off halfway through night 1), and 4 nights without playback. Here, 284 call sequences were recorded at the edge of the forest at playback site A, and 70 away from this edge in pasture: open site A. Because call characteristics were likely to differ depending on where these were made, we chose to compare only calls made at A’s edge site when there was and was not playback.

### Analysis

To assess the effect of playback on bat calls, we used generalised linear mixed effects models within the glmmTMB package (Brooks et al., 2017) in R 4.4.0 (R core team 2024), with separate models for the two bat species. The response variable was number of call sequences per night (counts). Playback was the fixed effect of interest. Detectors, nested within site, nested within paired site were included as random intercepts. We also included night as a random intercept, to account for differences in weather variation between nights. Count data were modelled with a negative binomial distribution with a linear parameterisation. The DHARMa package (Hartig, 2016) was used to check model diagnostics, including distribution assumptions, heteroscedasticity, and outliers. If there was a significant distribution violation in the ‘number of call sequences’ models, quadratic parameterisation of the negative binomial distribution was used. A dispersion parameter was included if tests indicated there was significant dispersion between playback and/or site parameters. For each model, a post-hoc contrast of playback was assessed using the emmeans package (Lenth, 2022), and results were plotted using the ggeffects package (Lüdecke, 2018).

Before testing the effect of playback, we assessed whether there was any carryover of the playback effect into the night after the playback, using the same model formats as described above, but using only those data from nights after playback occurred (e.g., playback occurred on the evening/night of the 9th-10th of February, so the evening of 10th-11th was tested for carryover effects, comparing sites with playback and sites without). No significant carryover effects were found (*p* = 0.29 for long-tailed bat; 0.06 for short-tailed bat), hence the nights after playback were included as controls in the main analyses that assessed the effect of playback.

To determine the effect of distance from the playbacks on bat call sequence detection we compared call sequence counts at the close detectors to those of the 30 m and 80 m detectors. For these analyses, data from two of the close detectors at each site were averaged to serve as values for comparison to the detectors placed into the forest, while the other two close detectors were averaged for comparison to the detectors placed along the forest edge (See Figure 2). Due to one faulty detector at Site C, only one close detector was used for the comparison against detectors into the forest for this site.

We found no difference in detection rates between the 30 m and 80 m detectors for nights with and without playback (*p* = >0.9 for all pairwise comparisons between 30 and 80 m detectors regardless of playback or bat species). This was likely due to the exponential decay of sound leading to similar exposure levels, detailed further in the discussion. We therefore present data as averaged (mean) to compare between close detectors and distant detectors, separately for forest and edge. As above, we used generalised linear mixed effects models separately for the two bat species. The models were the same, except for the addition of distance (either close or far) and its interaction with playback as fixed effects. Again, count data were modelled with a negative binomial distribution with a linear parameterisation. For each model, a post-hoc contrast of playback and distance was assessed using the emmeans package whenever there were significant interactions between predictors. P values were adjusted using the Holm-Bonferroni method (Holm, 1979).

## RESULTS

The number of bat call sequences varied between sites and throughout the study period (Figure 3). There was approximately an order of magnitude more long-tailed bat sequences recorded than short-tailed bat sequences (Figure 3).

**Figure 3.**
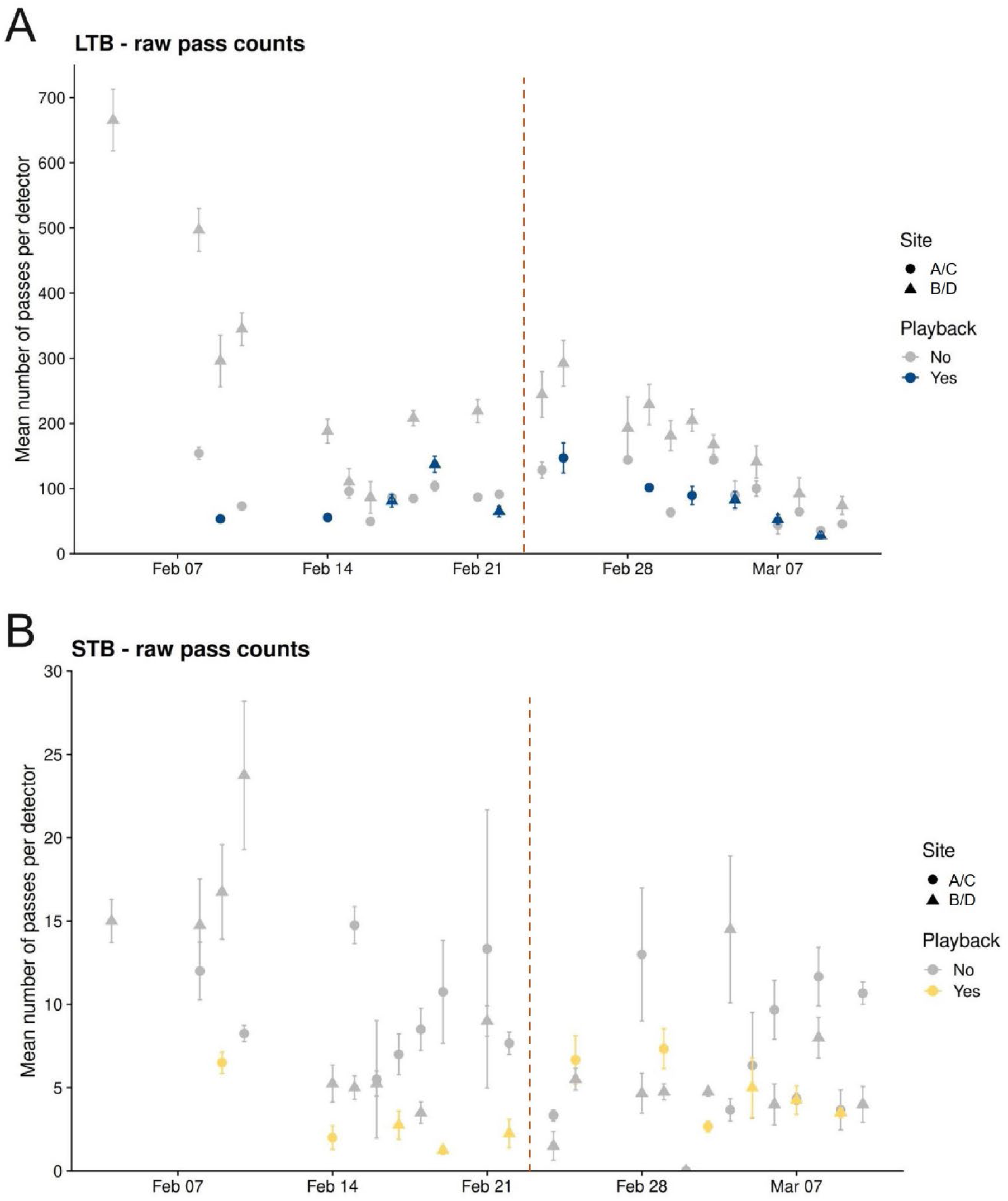
Mean number of A) long-tailed bat (‘LTB’, *Chalinolobus tuberculatus*), and B) lesser short-tailed bat (‘STB’, *Mystacina tuberculata rhyacobia*) sequences recorded per bat detector at Pureora Forest Park during playback time of recorded traffic sound for each night of the experiment (data shown as mean +/- SE; raw data is averaged from four detectors per site, apart from for Site C where it was from three detectors). Site pair 1 on Feb 5 was excluded due to part-night playback. The transition from Site Pair 1 to Site Pair 2 is indicated by the vertical red dashed line: during Feb 4 – Feb 22 the paired sites A & B were used, while from Feb 24 – March 10 paired sites C & D were used.

For long-tailed bats, there was a significant reduction in recorded sequences overall, as well as foraging sequences, during nights with playback compared with nights without playback (Figures 4A & 4B; Table 1). The effect of playback on the overall number of sequences recorded differed between sites, with no significant reduction recorded for Site C.

**Figure 4.**
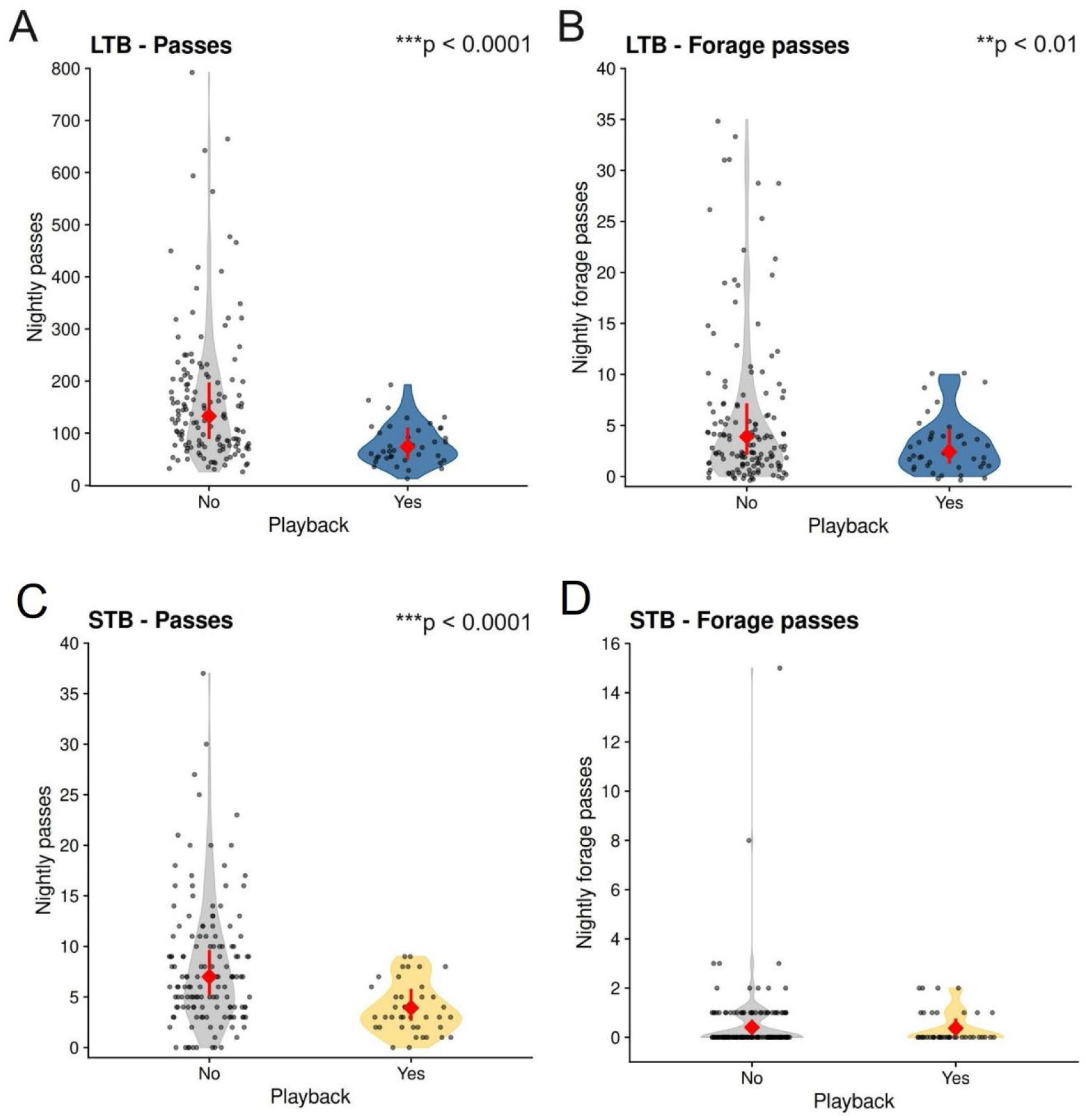
Modelled mean number of overall bat pass sequences (A,C) and foraging sequences (B,D), during nights with and without playback of traffic sound at Pureora Forest Park, for long-tailed bats (A-B, ‘LTB’, *Chalinolobus tuberculatus*) and lesser short-tailed bats (C-D, ‘STB’, *Mystacina tuberculata rhyacobia*). Grey points are raw data, red points and lines indicate back-transformed estimated marginal means with 95% confidence intervals, with coloured violin plots showing overall data distribution. P-values show significant differences for playback vs non-playback pairwise tests.

**Table 1.**
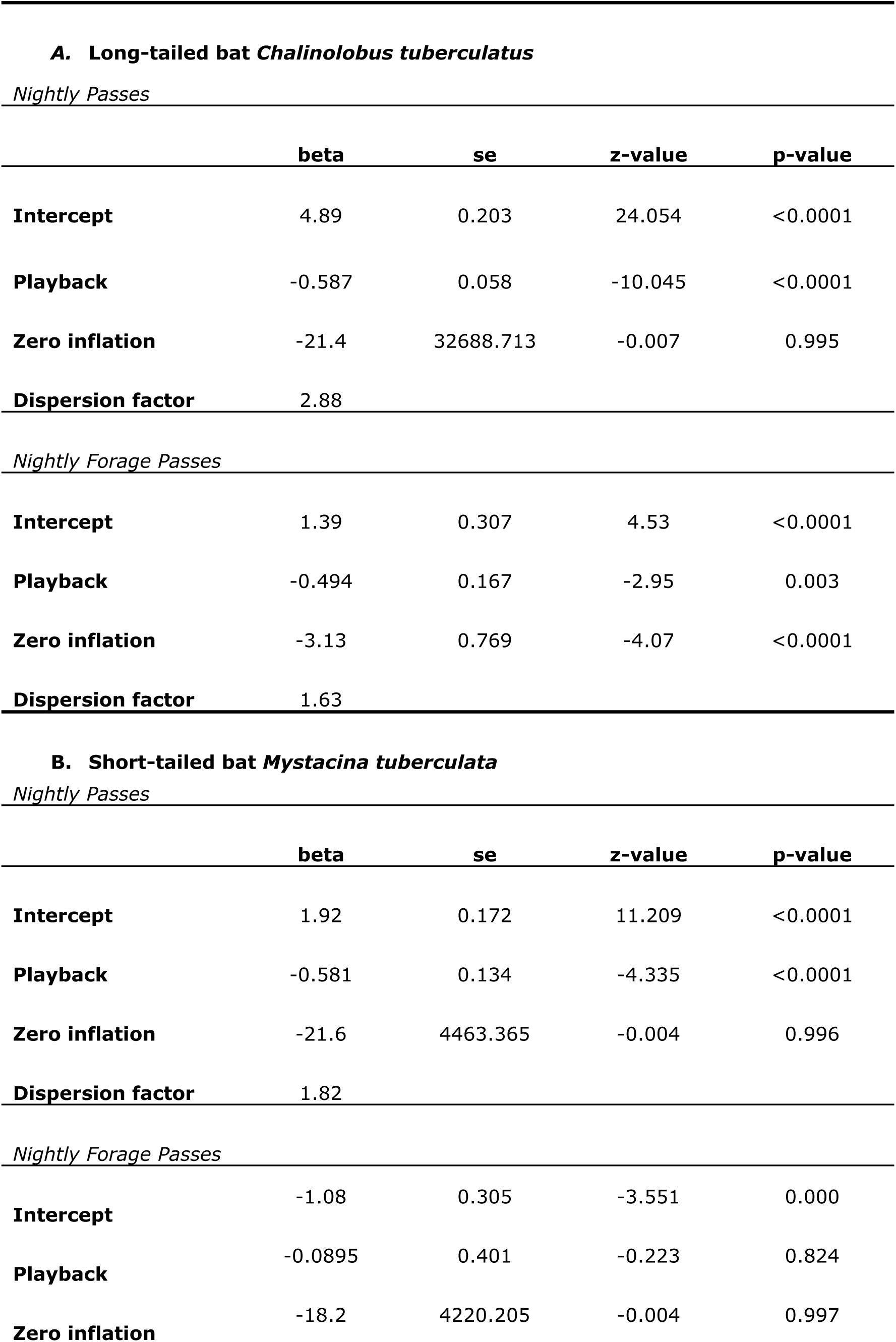
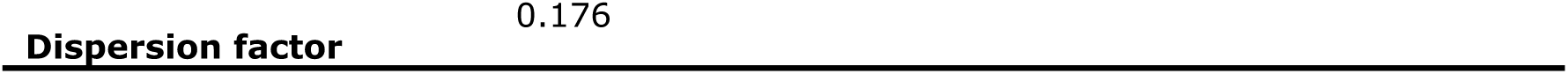
Results of negative binomial models of activity of long-tailed bats (A, *Chalinolobus tuberculatus*) and short-tailed bats (B, *Mystacina tuberculata*) at Pureora Forest Park, New Zealand, in response to playback of traffic sound.

The number of overall sequences of short-tailed bats also differed between playback and non-playback nights (Figure 4C), but foraging sequences did not (Figure 4D).

### Effect of distance from speaker on activity

We next examined whether the effects of playback on bat activity were dependent on distance from the speakers along the edge of the forest or into the forest. We found no difference in detection rates between the 30 m and 80 m detectors regardless of playback, and so used averaged data to compare between close detectors and distant detectors, separately for forest and edge. We found activity level at the distant detectors for either species did not differ based on whether playback was on or off, either along the forest edge or into the forest (Figure 5; Table 2).

**Figure 5.**
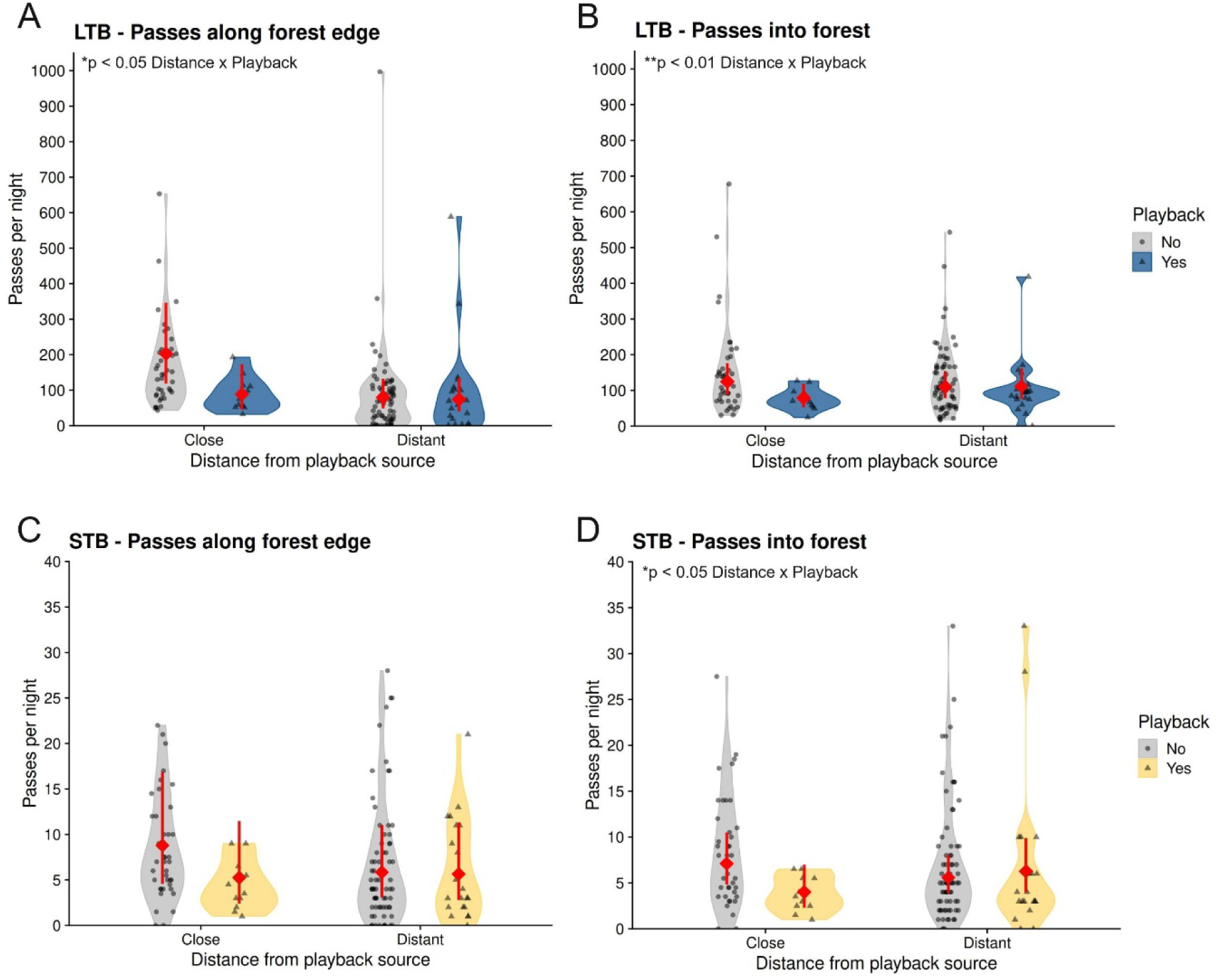
Average number of pass sequences for long-tailed bats (A & B, ‘LTB’, *Chalinolobus tuberculatus*) and lesser short-tailed bats (C & D, ‘STB’, *Mystacina tuberculata rhyacobia*), during nights with and without playback, as a function of distance from playback and orientation from forest edge. Distant = averaged 30 and 80 m detectors, Close = mean call sequence detections at two of the four close detectors, each conducted for comparisons into the forest and along the forest edge. Gray violin plots show the data distributions for nights when no playback occurred, coloured violin plots are for nights when playback occurred. Grey dots are the raw data and the red dot and lines represent means and 95% confidence intervals. Significant interactions between Distance and Playback are indicated where present: Playback was shown to affect bat activity at recorded on close detectors at sites A, B, and D, but not at distant detectors. Activity at distant detectors was not significantly different with vs. without playback for either bat species and in either distant orientation (into forest or along edge).

**Table 2.** Results of negative binomial models of activity of long-tailed bats (A, *Chalinolobus tuberculatus*) and short-tailed bats (B, *Mystacina tuberculata*) at Pureora Forest Park, New Zealand, at close and distance sites in response to playback of traffic sound.

| **A. Long-tailed bat *Chalinolobus tuberculatus*** |  |  |  |  |
| --- | --- | --- | --- | --- |
| *Nightly passes by distance along edge* |  |  |  |  |
|  | **beta** | **se** | **z-value** | **p-value** |
| **Intercept** | 5.31 | 0.274 | 19.39 | <0.0001 |
| **Playback** | -0.829 | 0.288 | -2.87 | 0.004 |
| **Distance** | -0.929 | 0.182 | -5.1 | <0.0001 |
| **Playback\*Distance** | 0.75 | 0.359 | 2.09 | 0.037 |
| **Zero inflation** | -2.72 | 0.357 | -7.63 | <0.0001 |
| **Dispersion factor** | 0.446 |  |  |  |
| *Nightly passes by distance into forest* |  |  |  |  |
| **Intercept** | 4.83 | 0.177 | 27.25 | <0.0001 |
| **Playback** | -0.454 | 0.139 | -3.27 | 0.001 |
| **Distance** | -0.127 | 0.075 | -1.7 | 0.090 |
| **Playback\*Distance** | 0.467 | 0.162 | 2.88 | 0.004 |
| **Zero inflation** | -4.994 | 1.004 | -4.92 | <0.0001 |
| **Dispersion factor** | 2.08 |  |  |  |
| **B. Short-tailed bat *Mystacina tuberculata*** |  |  |  |  |
| *Nightly passes by distance along edge* |  |  |  |  |
|  | **beta** | **se** | **z-value** | **p-value** |
| **Intercept** | 2.17 | 0.333 | 6.516 | <0.0001 |
| **Playback** | -0.514 | 0.275 | -1.87 | 0.061 |
| **Distance** | -0.407 | 0.155 | -2.625 | 0.009 |
| **Playback\*Distance** | 0.48 | 0.337 | 1.426 | 0.154 |
| **Zero inflation** | -1.998 | 5261.05 | -0.004 | 0.997 |
| **Dispersion factor** | 0.875 |  |  |  |
| *Nightly passes by distance into forest* |  |  |  |  |
| **Intercept** | 1.96 | 0.198 | 9.905 | <0.0001 |
| **Playback** | -0.572 | 0.251 | -2.278 | 0.023 |
| **Distance** | -0.242 | 0.122 | -1.983 | 0.047 |
| **Playback\*Distance** | 0.689 | 0.286 | 2.414 | 0.016 |
| **Zero inflation** | -20.08 | 5107.593 | -0.004 | 0.997 |
| **Dispersion factor** | 1.53 |  |  |  |

### Sound playback effects on call structure

Characteristics of long-tailed bat call sequences differed significantly during playback compared with other times (Figure 6; Table 3), despite few data points during playback. Call intensity (dB) was higher (less negative) and calls were shorter in length during playback than at other times (Figure 6A-B). Call sequences also had higher minimum, mean, and maximum frequencies and a higher mean bandwidth (difference between minimum frequency and frequency at the peak of the call) during playback (Figure 6C-E). Overall, this exploration suggests that bats call may be shorter, at a higher pitch and louder during playback compared with other times.

**Figure 6.**
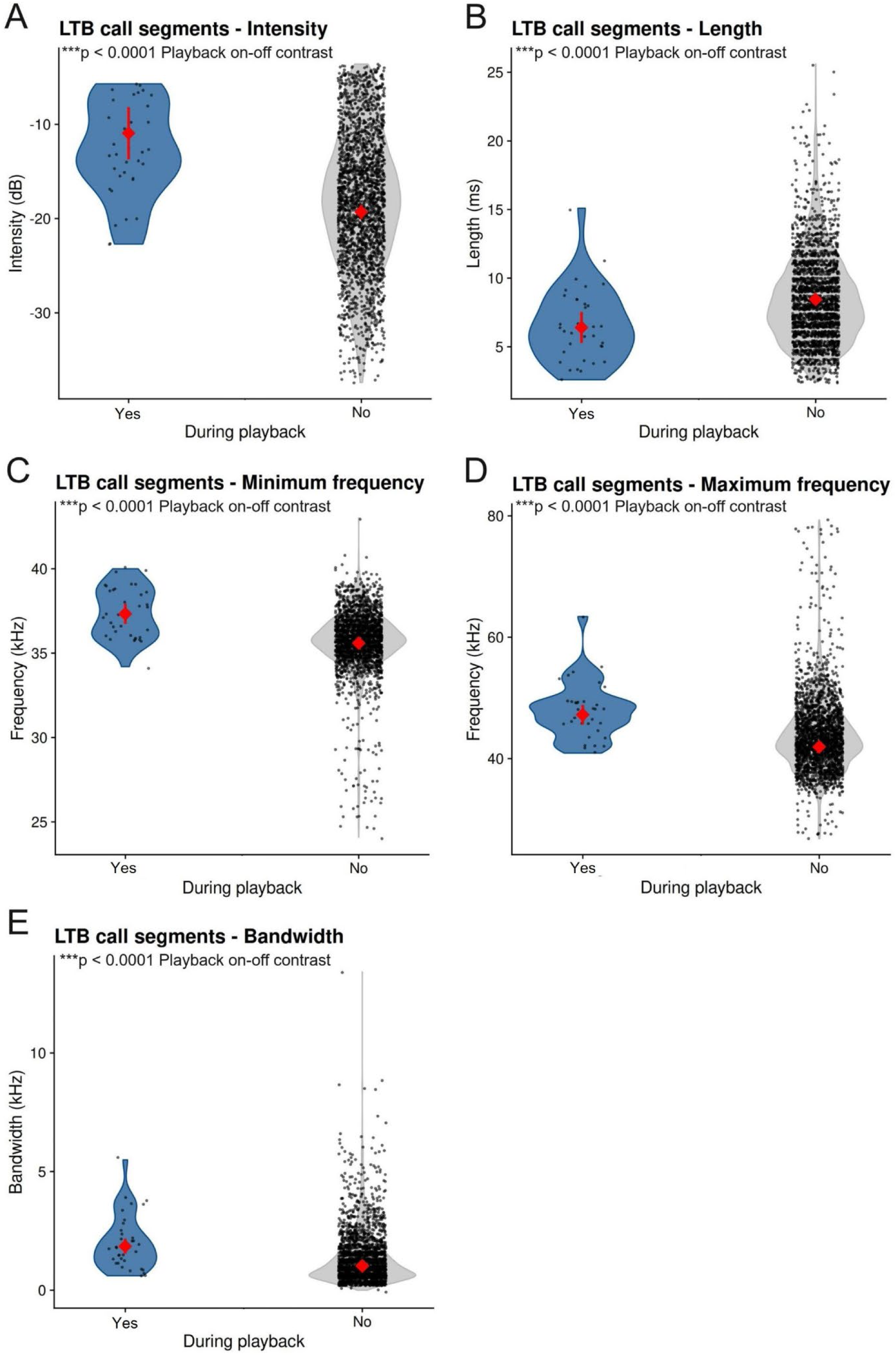
Characteristics of individual call segments during nights with and without playback, including intensity in dB (A), call length (B), minimum and maximum frequency (C & D), and bandwidth (E) of long-tailed bat (‘LTB’, *Chalinolobus tuberculatus*) call sequences at Site A during the 6 nights with information on individual call parameters (including 1.5 nights with playback). Grey violin plots show the data distributions for nights when no playback occurred, coloured violin plots are for nights when playback occurred. Black dots are the raw data and the red dot represent means.

**Table 3.** Results of t-distribution model contrasts of call characteristics of long-tailed bats (*Chalinolobus tuberculatus*) at Pureora Forest Park, New Zealand, in response to playback of traffic sound.

| *Call intensity* |  |  |  |  |
| --- | --- | --- | --- | --- |
|  | **beta** | **se** | **z-value** | **p-value** |
| **Intercept** | -11 | 0.415 | -26.5 | <0.0001 |
| **Playback** | -8.37 | 0.407 | -20.5 | <0.0001 |
| *Call length* |  |  |  |  |
| **Intercept** | 6.38 | 0.577 | 11.06 | <0.0001 |
| **Playback** | 2.04 | 0.561 | 3.64 | <0.001 |
| *Minimum frequency* |  |  |  |  |
| **Intercept** | 37.3 | 0.304 | 122.95 | <0.0001 |
| **Playback** | -1.74 | 0.284 | -6.11 | <0.0001 |
| *Maximum frequency* |  |  |  |  |
| **Intercept** | 47.2 | 0.828 | 57 | <0.0001 |
| **Playback** | -5.3 | 0.77 | -6.89 | <0.0001 |
| *Bandwidth* |  |  |  |  |
| **Intercept** | 1.85 | 0.163 | 11.34 | <0.0001 |
| **Playback** | -8.29 | 0.162 | -5.12 | <0.0001 |

## DISCUSSION

Here we used a phantom road design to determine whether bats exposed to the sounds from road traffic alter their behaviour by avoiding noisy areas or modulating their calls.

Along the phantom road, we found an approximate 50% reduction in the mean number of sequences per night of both long-tailed and short-tailed bats during sound playback, suggesting that bats are strongly avoiding the sound source within 30-50 m of the playback (the detection range of the recorders S. Cockburn as cited in Smith et al., 2020). The pronounced decrease in activity is unlikely to be due to decreased detection as the bat calls did not overlap spectrally with the playback. For long-tailed bats, the number of foraging call sequences decreased in line with overall call sequences, indicating this change was likely due to avoidance of the area, rather than the action of an independent mechanism. The number of short-tailed bat foraging call sequences were not significantly different during playback despite reduced overall activity; this may indicate a proportional increase in aerial hawking rather than gleaning/ listening for prey, or may simply be a product of the limited foraging data for this species. Our small dataset on call characteristics suggests that when long-tailed bats were present during traffic playback, their calls were shorter and louder, had greater bandwidth, and higher frequencies. Such frequency shifts are known from other frequency modulating echolocating bats when there is likely interference in their calls (Hiryu et al., 2010).

Both avoidance and call modulation are behavioural changes that may result in increased energetic costs for individual bats with flow-on effects for reproductive success (Kerth & Melber, 2009), fitness, and survival (Jones et al., 2019). We found no significant effects of sound playback on call sequence detection at the more distant locations (30 and 80 m), from the sound source, suggesting at this distance bats were not altering their activity level. We are aware that the majority of change in traffic noise levels occurs in the first 50 m, with little variation beyond 100 m (Berthinussen & Altringham, 2012), which may help explain this.

Avoidance behaviour in response to anthropogenic noise playback has been observed in a number of bat species (e.g., Finch et al., 2020; Luo et al., 2015; Schaub et al., 2008). However, there are few experimental examples - most research has been done with captive bats that do not have the option of avoidance (e.g., Gomes & Goerlitz, 2020; Jiang et al., 2019; Siemers & Schaub, 2011). In these cases, a decline in foraging efficiency or alteration of call characteristics has been consistently reported (e.g., Gomes & Goerlitz, 2020; Jiang et al., 2019; Siemers & Schaub, 2011). In the few captive studies where foraging bats had the option, avoidance was also observed (Luo, Siemers, et al., 2015; Schaub et al., 2008). Elsewhere, free-flying long-tailed bats were less likely to be detected during playback of aircraft noise compared to silent tracks, while there was little evidence for a decline in activity during overhead aircraft sequences (Le Roux, 2010; Le Roux & Waas, 2012). In the four other experimental studies we are aware of that involve anthropogenic noise playback with wild, free-living bats, avoidance behaviour was observed across three of them (Domer et al., 2021; Finch et al., 2020; Hooker et al., 2023), although not in all bat species present (Hooker et al., 2023). One traffic playback experiment found a marked increase in activity for *Noctilio albiventris* during playback, theorised to be due to an increase in foraging effort required in noisy conditions; this was supported by alteration of several call characteristics (Yantén et al., 2022). These experiments with wild populations generally echo correlative findings around existing sources of anthropogenic noise (e.g. Bunkley et al., 2015; Wang et al., 2022). However, the majority of these studies had limited power to explore the mechanisms driving the avoidance behaviour. A recent meta-analysis of the effects of anthropogenic noise on bat behaviour found that habitat preference aids in explaining the differing results between species (Li et al., 2025).

However, the general agreement of research findings from studies with captive and free-living bats, as well as the results presented here, suggests that avoidance of anthropogenic noise is a widespread (although not ubiquitous) response amongst bats that use laryngeal echolocation. The range of species’ traits and behaviours such as call structure, flight speed, habitat use, and foraging strategies represented in these findings are broad and continue to grow (Finch et al., 2020). Given the apparent tendency for avoidance across the functional groups of laryngeal echolocating bats studied so far, broader research should be considered to assess potential effects of anthropogenic noise on other echolocating bats (i.e. lingually echolocating bats in the genus *Rousettus*) and non-echolocating bats (Pteropodidae, excepting *Rousettus*) that are generally considered to have lower acoustic reliance (Berkovitz & Shellis, 2018), in addition to other fauna overlooked in noise studies for similar reasons. It should be noted that this variation in species traits means that the mechanisms driving avoidance behaviour in bats may differ between species, or vary in relative importance in cases where avoidance is caused by multiple factors (e.g., masking and distraction). There may additionally be significant variation between individuals or changes in effect over time (Gomes & Goerlitz, 2020).

Our study did not explicitly assess the mechanism(s) that drove the reduction in bat activity near the phantom road. Although the playback included ultrasonic frequencies, anthropogenic noise such as the traffic recording used is low frequency and no overlap was apparent on the recordings produced by either detector type. However, it has recently been theorised that masking can occur with non-spectrally overlapping noise due to groupings of bat neurons that respond to background noise; the neuronal activity associated with processing the playback may therefore overlap and interfere with processing of higher-frequency signals (Zhang et al., 2023; Zou et al., 2023).

We acknowledge that data collection with the full-spectrum recording detectors was limited during playback, and changes in intensity are challenging to measure in the field as they are impacted by factors such as environmental conditions, clutter, and distance of the animal from the microphone (Attenborough, 2014; Pater et al., 2009). However, if there was no change in call loudness, then avoidance of the playback source would be assumed, if anything, to lead to lower average intensities measured, rather than higher as we observed. Our explorative findings of long-tailed bats calling at higher intensities and frequencies in the presence of anthropogenic sound is consistent with research in other bat species (e.g., Gomes & Goerlitz, 2020; Luo, Goerlitz, et al., 2015; Pedersen et al., 2024). A key concept to consider is the signal-to-noise ratio (SNR), which is a measure of the strength of the bat call or resulting echo (signal) to the irrelevant background noise (sound playback) (Corcoran & Moss, 2017). A higher SNR makes the signal easier to detect within the background noise. The Lombard effect describes the increase in vocal intensity in noisy environments that aids in maintaining an appropriate SNR and has been well-documented in many vertebrates (Kunc et al., 2022). This effect is often considered to be involuntary, such as a person speaking loudly when they are wearing headphones, and may also result in other vocal characteristics, like bandwidth and frequency, being altered (Hotchkin & Parks, 2013). The Lombard effect may be linked to both masking and distraction (Allen et al., 2021; Pedersen et al., 2024).

Other than a by-product of the Lombard effect, frequency shifting is often used to avoid masking. This can be seen in many bat species both in playback experiments (e.g., Bates et al., 2008; Hage et al., 2013) and in natural vocalisations, such as where bats in proximity appear to adjust their call frequencies to avoid overlap with each other (Ulanovsky et al., 2004). Further experimental research is needed to untangle the mechanisms driving the observed changes in call characteristics. Interestingly, Finch et al. (2020), who carried out the first published experimental research to assess the response of free-living bats to traffic noise, found a significant reduction in activity for all five bat species in proximity to their phantom road during playback of the non-ultrasonic component. However, an effect was only observed in one of the species when just the ultrasonic component was played. This could be due to several reasons: limited sound in the ultrasonic frequencies to drive an effect, the rapid attenuation of higher frequency sounds compared to lower, or masking simply not being the primary driver of avoidance for the species present.

Short-tailed bats may be particularly sensitive to masking or distracting noise, as they spend a large proportion of their foraging time gleaning from the forest floor, relying largely on adventitious sounds from invertebrate prey (Jones et al., 2003). These cues (i.e. prey rustling in leaf litter) often include lower frequency components that have notable spectral overlap with anthropogenic noise compared to typical ultrasonic bat vocalisations (Goerlitz et al., 2008; Schaub et al., 2008; Siemers & Schaub, 2011). In this scenario, bats would be unable to mitigate decreased detection through call modulation. This means likely effects for short-tailed bats are either decreased foraging efficiency (Bunkley & Barber, 2015), avoidance of noisy areas (Schaub et al., 2008), or preferentially employing aerial hawking of prey rather than substrate gleaning (Gomes et al., 2021). All may result in a reluctance in short-tailed bats and other similar gleaning bats to stay near a noisy road or to cross it (e.g., *Myotis bechsteinii*; Kerth & Melber, 2009). These barrier effects may be more powerful than for bats with other preferred feeding methods and wing morphologies (Kerth & Melber, 2009).We found short-tailed bat detection rates declined during playback, but the number of short-tailed bats’ foraging call sequences did not differ depending on whether there was playback. This suggests short-tailed bats were increasing the rate at which they aerially hawked for prey, whilst also avoiding the area during playback. Whether this represents a change in foraging strategy that persists when sites are noisy over longer periods requires further research.

Key questions that remain are what the threshold is for the effects we have observed, and whether bats habituate to noisy environments. While we found evidence of an avoidance effect close to the sound source (with detectors placed close to the sound source) reflective of bats flying 55 m from the phantom road, we did not at 30 m and 80 m from the phantom road. This could represent bats echolocating 80-130 m from the phantom road given known detection distances of the recorders used. However, this distance is not comparable to sound levels 30 or 80 m from an actual highway. Although the sound pressure levels along our phantom road (∼73 dB) were comparable at source to a busy road (Jackett, 2023; Wilkening, 2012), the resulting soundscape differed in several regards, as is often the case for artificially simulated noise (Pater et al., 2009). The speaker array used three groups of speakers spaced over a relatively short distance, and the sound was directional and attenuated rapidly. When 30 m into the forest interior, sound levels during playback averaged 43 dB and at some sites, heavy vehicles from the state highway >500 m away could occasionally be heard over the sound playback. A meta-analysis of terrestrial fauna responses to anthropogenic noise found that effects begin to appear at around 40 dBA (Shannon et al., 2016). The AR4 detectors used also have a range of 30 – 50 m, each covering a broad range of sound pressure levels during playback trials and overlapping with more distant detection points. If anything, we suspect these factors are likely to have led to an underestimation of effects compared to real noise pollution from a physical highway i.e., we suspect that the effect of noise from a real road would be greater and may persist over a larger area. When long-tailed bat detection rates were compared between sites alongside highways and similar sites away from highways, higher rates of detections were found at sites 200 m away from highways (Borkin et al., 2019). Together with our research, this suggests that the effect of actual highway traffic noise on bat activity may be reduced as far away as 130 to 200 m from the source.

Because many environmental factors influence sound dispersion (temperature, humidity, wind, etc.; Attenborough, 2014), future studies should measure the ‘effective exposure’ of sound at each detector for every night of playback. Such studies would be valuable for determining the maximum distance range at which bats are affected by traffic noise, although this range is likely highly variable depending on road design, frequency of use, and the surrounding environment.

Further study is required to untangle what mechanisms are driving the observed call modulation and avoidance, and whether key behaviours are inhibited for individuals who do not avoid noisy areas. Call modulation may be an energetically costly strategy with resultant fitness effects (Currie et al., 2020). Avoidance of an area is effectively habitat loss; chronically noisy environments may effectively exclude bats from key resource locations such as of prey and roost trees. Line noise sources such as busy roads likely present a barrier to movement and increase habitat fragmentation, whilst quieter, lower traffic volume roads act more as filters to movement (Bennett & Zurcher, 2013). Indeed, rates of detection of long-tailed bats are lower alongside major roads compared to similar habitat nearby (Borkin et al., 2019). Our research, and that of Borkin et al. (2019) and Schamhart et al. (2024) suggests a combination of noise, light, and traffic volumes may influence the scale of effect of roads – and the Zone of Influence. The effect may increase further with increasing traffic volumes (Borkin et al 2019). Research into whether bats habituate to such noisy environments and their tolerance for different sources of anthropogenic noise is imperative to understand how we can manage impacts of future development on sensitive fauna.

### Conclusion & implications

In summary, we found evidence for a decline in the overall activity of long-tailed bats and short-tailed bats, and a decline in long-tailed bat foraging activity, in response to playback of traffic sound. A ∼50% decrease in estimated mean sequence number indicates bats were avoiding the area during playback, although this effect was not apparent at distant detection points. Long-tailed bats also appeared to alter their calls during playback to call more loudly and at higher frequencies despite no spectral overlap between the sound playback and their typical calls. These strategies may impose fitness costs on bats through increased energy expenditure or by high sound levels acting as an invisible barrier to resources and reducing quality habitat. Evidence to support noise carryover effects was generally weak or non-existent, and more targeted research is suggested in this area, in addition to looking at effects of high sound levels on roost emergence.

With increasing areas of natural habitat exposed to anthropogenic noise, untangling the effects of high sound levels is therefore crucial for the protection of bats and the ecosystem services they provide.

## Author contributions

KB, TP, DP, and AH conceived and designed the study. AH, KB, TP, and DP conducted the experiment and collected the data. EC and PM performed the analyses, with figures prepared by EC, PM, DP, and AH. AH, EC, KC, KB, DP, and PM contributed to writing the manuscript. DP, KB, and KC provided project supervision. KB secured Animal Ethics approval and arranged funding from the Department of Conservation Bat Recovery Group and Department of Conservation Threatened Species Research Workstream. DP arranged support from Plant and Food Research through vehicle and fuel provision during data collection. AH was supported by a University of Auckland Master’s scholarship covering tuition and a living expenses stipend during the research.

